# Hippocampus mediates conceptual generalization of pain modulation

**DOI:** 10.64898/2026.10.16.759532

**Authors:** Dylan Sutterlin-Guindon, Tor D. Wager, Leonie Koban

## Abstract

Pain is strongly influenced by expectations and learning from previous experience, such as in classical conditioning. Conditioned responses and expectations can generalize to perceptually and conceptually related cues, but how generalization influences pain experience and the neurobiological processing of pain remains unclear. We used fMRI and multilevel mediation analyses to address this question. Thirty-six human participants first learned to associate two visual cues from distinct conceptual categories (e.g., animals vs. vehicles) with high or low levels of heat pain. In a subsequent phase, they were presented with novel cues (images, drawings, or words) not previously paired with pain, but which shared the conceptual category of the initial pain-predictive cues. Participants who developed explicit expectations during learning reported greater pain in response to stimuli conceptually related to high-vs. low-pain cues (‘generalization stimuli’), demonstrating generalization of cue influences on pain. This effect was mediated by increased pain-related activity to generalization stimuli in the hippocampus, which correlated with individual differences in cue-evoked expectations. A broader network, including areas of the default mode network and striatum, also contributed to conceptual generalization of pain modulation, while threat-related regions such as the amygdala responded to generalization stimuli but did not mediate effects on pain ratings. These findings extend our understanding of expectancy-driven pain modulation by showing how conceptual processes can influence pain and its neurobiological substrates, offering new insight into placebo effects and maladaptive learning in chronic pain.

## Introduction

Pain is the leading reason people seek medical care, imposing substantial costs on health systems and individual well-being (Jackson et al., 2016). However, pain is not merely a response to tissue damage. Modern conceptions of pain suggest that it is an inference constructed in the brain about potential damage (Seymour, 2019), influenced by predictive processes—including placebo and nocebo effects (Büchel et al., 2014; Colloca, 2019; Kaptchuk et al., 2020; Wager & Atlas, 2015)—as well as nociceptive input. Pain, like other experiences and physiological processes related to harm and reward, can be influenced in advance by predictive cues, which are thought to prepare the organism for survival-relevant challenges and opportunities (Eikelboom & Stewart, 1982; Ramsay et al., 2020; Smith et al., 2021). In some cases, cue effects are mediated by expectations, suggesting that conscious, reportable processes are important (Jepma & Wager, 2015; Kirsch, 1985; Koban & Wager, 2016). In other cases, conditioned associations can occur outside of conscious awareness (Benedetti et al., 2003) and be independent of explicit expectations (Goebel et al., 2002; Schafer et al., 2015). This literature demonstrates multiple pathways by which the brain draws on experience to predict future pain. Identifying the neural mechanisms that support this predictive process is essential for improving treatment and reducing the burden of chronic pain (Motzkin et al., 2023). Here, we test the brain mechanisms of *conceptual generalization* of previous experiences on pain.

Critically, expectations often extend beyond specific prior experiences. Instead, people can generalize across conceptually related situations by relying on abstract similarities (Kumaran, 2012). In aversive learning, for example, fear generalizes to new situations based on their resemblance to previous threats (Dunsmoor & Murphy, 2015). Similar generalization processes may influence pain. People expect pain in novel settings that resemble painful ones (Glogan et al., 2023), and such mechanisms likely contribute to placebo and nocebo effects (Weng et al., 2025). For instance, expectations of relief from one drug may transfer to another medication if the two are perceived as serving the same purpose. Alternatively, if a treatment fails, it may create negative expectations that reduce the effectiveness of future treatments (Zunhammer et al., 2017). Yet, how the brain supports conceptual generalization in pain remains unknown.

Associative learning plays a central role in shaping how we experience pain. In classical conditioning paradigms, a neutral cue used as a conditioned stimulus (CS) is repeatedly paired with either high or low pain. Over time, the CS comes to signal expected pain intensity, and this learned association influences subsequent pain experiences. Cues previously paired with high-intensity (CS_HIGH_) compared to low-intensity noxious stimuli (CS_LOW_) lead to increased pain ratings, even when the stimulus intensity is kept constant across trials (Atlas & Wager, 2012; Jepma & Wager, 2015; Traxler et al., 2019; Zhang et al., 2019). These learned expectations induced by pain-predictive CSs not only shape subjective pain, but they also change how the brain responds to pain. Functional imaging studies have linked this pain modulation to increased pain-related activity in the thalamus, posterior insula, sensorimotor cortices, and anterior mid-cingulate cortex, as well as in frontal areas involved in attention, value, and expectation (Atlas et al., 2010, 2022; Jepma et al., 2018; Koban et al., 2019; Koyama et al., 2005; Necka et al., 2025). Other systems, including the hippocampus and medial prefrontal cortex, are also key targets for explaining how learned expectations influence pain (Bingel et al., 2022; Motzkin et al., 2021).

While associative learning explains how cues and contexts shape pain processing, it cannot account for how novel cues, never directly paired with pain, can influence pain perception. Generalization allows the predictive value of previously learned stimuli to transfer to new ones (Dunsmoor et al., 2011; Dunsmoor & Murphy, 2015; Honig & Urcuioli, 1981). Most research has focused on perceptual generalization, where generalization stimuli that resemble CSs in their visual or sensory features evoke similar reactions (Kampermann et al., 2021; Lissek et al., 2014; Liu et al., 2019). In contrast, conceptual generalization involves the transfer of learning across abstract or categorical dimensions, independent of perceptual similarity (Kumaran, 2012). Behavioral studies show that both (a) self-reported measures of fear, avoidance and pain expectancy and (b) physiological measures like eye-blink startle and skin conductance responses can generalize based on conceptual similarity to cues (Mertens 2021, Meulders 2017, Meulders & Bennet 2018; Kloos 2022). Also, these self-reported measures and physiological responses can generalize across various modalities, including conceptual categories, physical movements, and real-world representations (Dunsmoor & Murphy, 2015; Glogan et al., 2021; Kloos et al., 2022). In these cases, generalization stimuli never paired with pain can bias subsequent affective responses through expectations developed during initial learning, and these biases can be extinguished if the expected outcome does not follow (Glogan et al., 2019).

Despite these findings, the impact of conceptual generalization on pain perception itself has received less attention. In a series of three behavioral experiments, Koban et al. (2018) examined both perceptual and conceptual forms of pain generalization and consistently found that generalization effects on pain depended on individual differences in the strength of explicit learning and associated pain expectations: Participants who formed stronger expectations that a cue predicted high or low pain during the learning phase showed stronger pain modulation during generalization. Liu et al. (2019) also reported increased pain ratings following high versus low pain-related conceptual generalization cues. However, the brain mechanisms supporting this process and underlying the individual variability in learning and generalization effects on pain remain unknown. To address this gap, the present study investigates the neural mechanisms through which learned pain expectations generalize across conceptual categories to modulate pain responses.

Several candidate brain systems may mediate conceptual generalization effects on pain. On the one hand, generalization effects may rely on regions involved in threat detection and aversive learning, such as the amygdala, insula, anterior mid-cingulate, and striatum (Dunsmoor et al., 2011; Lissek et al., 2014). On the other hand, conceptual generalization effects on pain may depend more on brain regions involved in conceptual and semantic processing, such as the hippocampus and areas of the default mode network (DMN). Neuroimaging studies of perceptual generalization in fear conditioning show that both threat learning-related circuits and regions central for conceptual processing, such as the hippocampus and ventromedial prefrontal cortex (vmPFC), contribute to the reinstatement of learned fear responses (Dunsmoor & Murphy, 2015; Onat & Büchel, 2015; Webler et al., 2021, 2024). Across tasks and modalities, the hippocampus has been shown to support cognitive maps—structured representations that capture abstract relationships across spatial, conceptual, and social domains (Constantinescu et al., 2016; Garvert et al., 2017; Sherrill et al., 2023; Tavares et al., 2015). In interaction with key hubs of the DMN, such as the vmPFC, the hippocampus facilitates the integration and generalization of learned associations to novel situations by drawing on high-level features (Bowman & Zeithamova, 2018; Vatansever et al., 2017). This system is thus well-positioned to support the transfer of pain-predictive value from CSs to conceptually related stimuli, and bias pain.

Here, we used fMRI to identify the brain mechanisms underlying pain modulation induced by conceptual generalization. Thirty-six participants first learned to associate visual cues from different conceptual categories (e.g., animals vs. vehicles) with high or low levels of heat pain (CS_HIGH_ and CS_LOW_). In a subsequent generalization phase, they received medium-intensity heat stimulation following novel generalization stimuli (GS) presented in different modalities (images, drawings, or words). Although never paired with pain, these GS belonged to the same conceptual categories as either the high- or low-pain learning cues (GS_HIGH-CAT_ and GS_LOW-CAT_). We then used multilevel mediation analyses (Atlas et al., 2010; MacKinnon et al., 2007; Wager et al., 2009) to identify brain regions in which pain-evoked activity was modulated by conceptual generalization and explained trial-by-trial variation in pain ratings. We further tested the hypothesis that individual differences in expectations would moderate these effects, consistent with prior behavioral findings (Kloos et al., 2022; Koban et al., 2018).

## Results

### Behavioral results

During an associative learning procedure, participants learned to associate two conditioned stimuli (CS) with either high (CS_HIGH_) or low (CS_LOW_) levels of pain. To test how these cues influenced pain ratings independently of actual stimulation intensity, medium-intensity (49°C) painful stimuli were presented on 50% of trials within both CS_LOW_ and CS_HIGH_ conditions. Participants were not told which cues predicted high versus low pain levels. They were only instructed that one cue would predict high and the other cue low pain, and that they had to discover the cue-pain contingency. Results from this associative learning phase were previously reported (Koban et al., 2019). In brief, on medium-intensity trials, CS_HIGH_ led to higher pain ratings compared to CS_LOW_ (*β* = 0.42, 95%-CI: [0.05–0.79], *t*(35) = 3.13, *p* = 0.036, Cohen’s *d* = 0.52). Participants also reported their pain expectations before each stimulus (see Figure 1a). CS_HIGH_ led to higher expectations than CS_LOW_ on medium-heat trials (*β* = 1.41, 95%-CI [0.51–2.31], *t*(35) = 3.09, *p* = 0.004, *d* = 0.50). These results confirm that pain expectation and pain ratings were successfully altered by the pain-predictive cues (see Figure 2a-b), but with substantial individual variability in the strength of associative learning. In the present study, we focus on the subsequent conceptual generalization task (not previously reported), and how the learning strength (degree of explicit learning) during the associative learning task influenced subsequent generalization effects on pain.

**Figure 1.**
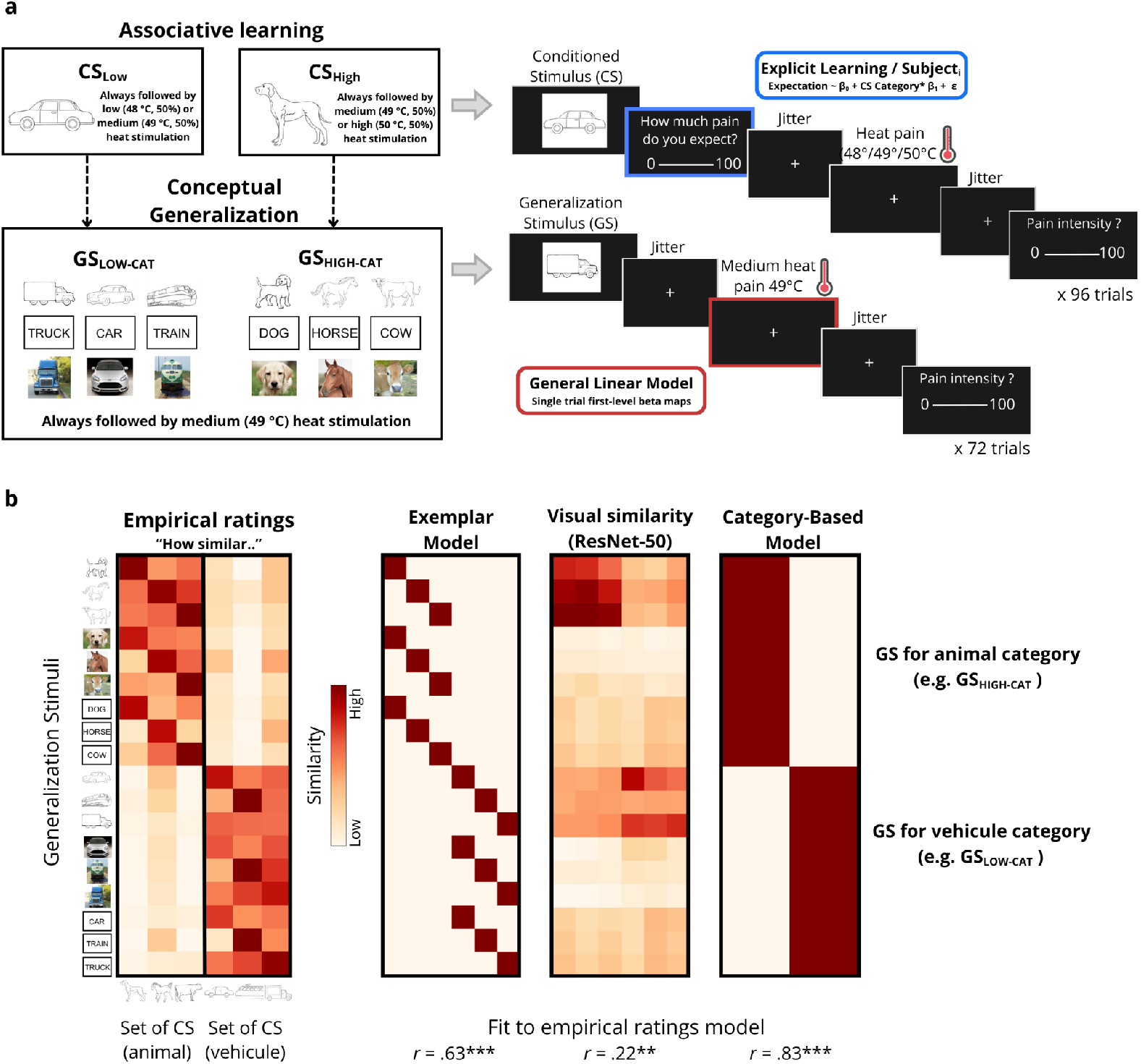
Paradigm and experimental stimuli. **a)** *Associative learning task* (previously reported in Koban et al., 2019). Participants learned to associate two conditioned stimuli (CS) with different levels of heat pain. The CS_LOW_ cue (e.g., a vehicle) was followed by either low (48 °C, 50% of trials) or medium (49 °C, 50% of trials) heat. The CS_HIGH_ cue was followed by either high (50 °C, 50% of trials) or medium (49 °C, 50% of trials) heat stimulation to the right calf. On each trial, participants first viewed a CS cue (at the same time as social ratings that were independent of the CS cues; see Koban, et al., 2019), rated expected pain (0-100 visual analog scale), received heat stimulation, and then rated pain (0-100 scale). Expectancy ratings from this task (blue box) were used to quantify individual differences in explicit learning of the cue-pain contingency, computed as the slope (*β_1_*) of expected pain ratings by CS category. *Conceptual generalization task*. In the subsequent generalization phase, participants viewed generalization stimuli, i.e., novel exemplars from the same conceptual categories as the CSs, that were never paired with heat stimulation during learning. The GS_HIGH-CAT_ stimuli (e.g., dog, horse, cow) belonged to the same category as CS_HIGH_, while the GS_LOW-CAT_ stimuli (e.g., truck, car, train) belonged to the same category as CS_LOW_. Three cue modalities were used during generalization: cartoon, picture, and word. All generalization stimuli were followed by medium heat stimulation (49 °C) without expectation rating. At the end of each trial, participants rated how much pain they experienced, again using a 0-100 VAS). Pain-related brain activity during this phase (red box) was modeled using a single-trial general linear model, producing trial-by-trial beta maps. These maps were then used in subsequent mediation analyses. **b)** Empirical ratings of similarity between conditioned and generalization stimuli compared with three different similarity models. Participants rated the similarity between each of the 18 generalization stimuli and each CS on a continuous scale, without specific instructions about whether to focus on visual or conceptual features. The first heatmap shows the empirical similarity matrix (median ratings across participants), where darker red indicates greater perceived similarity. We tested three models to characterize the structure of these ratings, with Spearman rank correlation and estimating significance with 10,000 permutations. The *exemplar model* assumed similarity only between identical concepts (e.g., dog CS = dog across all modalities), ignoring category structure. The *ResNet-50 model* was derived from a deep neural network (He et al., 2016) to quantify low-level visual similarity. The category-based model assumed maximal similarity among all stimuli from the same conceptual category (e.g., animals), regardless of exemplar or modality. The *category-based model* best explained the empirical ratings (rho = 0.83, *p* < 0.001), outperforming the exemplar (rho = 0.63, *p* < 0.001) and visual similarity (rho = 0.21, *p* = 0.01) models, supporting a categorical representation of generalization stimuli (GS_LOW-CAT_ and GS_HIGH-CAT_). \*\*\**p* < 0.0001, \*\**p* < 0.01

**Figure 2.**
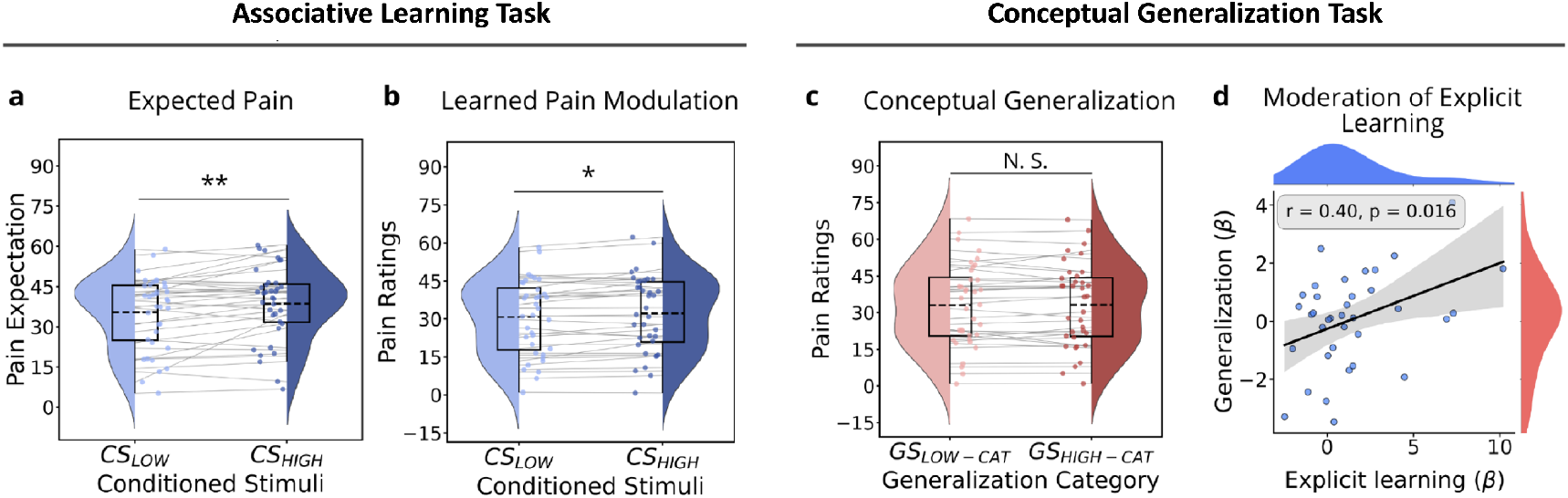
Behavioral effects. We used multilevel general linear models to characterize the behavioral effect of experimental conditions on pain expectancy and pain ratings for the associative learning task (previously reported in Koban et al., 2019) and for the generalization task. **a)** Expected pain during associative learning. Participants reported higher pain expectations for CS_HIGH_ versus CS_LOW_ stimuli during medium-intensity trials (49°C, 50% of trials). Explicit learning scores were quantified as the individual slope (*□_1_*) of the effect of CS category on expected pain and used in subsequent moderation analyses. **b)** Learned pain modulation during associative learning. Pain ratings were higher for CS_HIGH_ versus CS_LOW_ stimuli on medium-intensity trials. **c)** Pain ratings during conceptual generalization. Pain ratings did not significantly differ between GS_HIGH-CAT_ and GS_LOW-CAT_ conditions overall (n.s.). **d)** Moderation by explicit learning. Individual differences in explicit learning significantly moderated the strength of conceptual generalization effects on pain ratings during GS trials. This moderation is illustrated in the scatterplot by the positive association between individual generalization effect (*□*) and individual explicit learning scores (*□*). \**p* < 0.05; \*\**p* < 0.01

In the generalization phase, the same participants received medium-intensity pain stimulation following generalization stimuli (GS) that were never paired with pain but belonged to the same conceptual category as either the high- or low-pain CSs. At the group level, conceptual generalization (GS_HIGH-CAT_ > GS_LOW-CAT_) did not significantly influence pain ratings (*β* = 0.12, 95%-CI: [-0.35–0.59], *t*(35) = 0.46, *p* = 0.59, *d* = 0.04). However, consistent with previous studies (Koban et al., 2018), participants who developed more explicit knowledge of the CS-pain contingency (as reflected in higher expectation ratings for CS_HIGH_ than CS_LOW_) showed larger generalization effects on pain, as revealed by a moderation of generalization effects by individual differences in pain expectation during the associative learning phase ([GS_HIGH-CAT_ > GS_LOW-CAT_] x expectancy interaction, *β* = 0.24, 95%-CI: [-0.01–0.49], *t*(35) = 2.46, *p* = 0.029, *d* = 0.41). Non-parametric permutation tests further confirmed the significance of this moderation effect (5,000 perm., *p* = .0016). These findings indicate that participants who became aware of the CS-pain contingency reported higher pain ratings during GS_HIGH-CAT_ compared to GS_LOW-CAT_ trials, showing generalization effects on pain (Figure 2c-d).

### Brain mediators of the conceptual generalization effect on pain

To investigate the brain mechanisms underlying conceptual generalization, we conducted a mediation effect parametric mapping analysis (Figure 3), testing for within-person mediation of cue effects on pain and moderation by individual differences in expectation ratings in each brain voxel (Atlas et al., 2010; Wager et al., 2008). The multilevel mediation used trial-by-trial cue values (GS_HIGH-CAT_ or GS_LOW-CAT_) as the initial variable, trial-by-trial brain activity as the mediator, and trial-by-trial pain reports as the outcome at the first (within-person) level of the model. *Path a* reflects cue effects on the brain, *Path b reflects* brain effects on pain, and the mediation effect *ab* reflects their joint strength. The second level tests the significance of each path (*a*, *b*, and *ab*) with participant as a random effect, along with moderation of each path by individual differences in explicit learning (average difference in pain expectation for CS_HIGH_ - CS_LOW_) as a second-level moderator (i.e., moderated mediation; see Figure 3). We report significant brain regions for each path (*a*, *b*, and *ab*) FDR corrected at *q* < .05. Full statistical results, including peak coordinates, Z-values, and cluster sizes, are provided in Supplementary Tables 1-3. *Path a* tested whether brain activity during pain differed as a function of generalization conditions (GS_HIGH-CAT_ > GS_LOW-CAT_). This contrast revealed the strongest activations (*q* < 0.05 FDR corrected) in the bilateral amygdala and in the left mid-posterior insula (Figure 4a). Additional regions showing increased activation included the right premotor cortex, right primary motor cortex, left posterior hippocampus, and subgenual anterior cingulate cortex (sgACC). These findings indicate that GS_HIGH-CAT_ stimuli (i.e., conceptually related to CS_HIGH_) elicited stronger pain-related brain responses in these regions. In contrast, a small cluster in the right dorsolateral prefrontal cortex (dlPFC) showed a negative effect, indicating reduced activation during GS_HIGH-CAT_ > GS_LOW-CAT_ trials.

**Figure 3.**
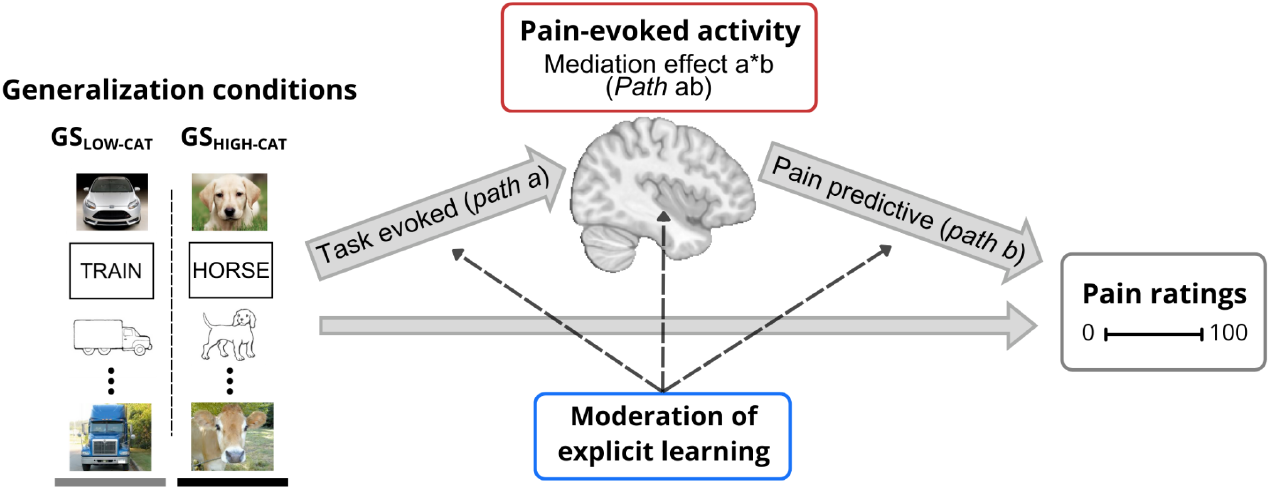
Multilevel moderated mediation of pain-evoked brain activity. We conducted a multilevel mediation effect parametric mapping analysis to identify brain regions involved in conceptual generalization effects on pain processing and pain ratings. Trial-by-trial pain-evoked brain activity was used to test: (1) brain regions showing greater activation during GS_HIGH-CAT_ > GS_LOW-CAT_ trials (*Path a*), (2) brain activity associated with higher pain ratings on a trial-by-trial basis, controlling for *path a* effects (*Path b*), and (3) brain regions formally mediating the effect of conceptual generalization condition on pain ratings (*Path ab*). We also tested whether the mediation effect (*Path ab*) was moderated by individual differences in explicit learning of the initial CS-pain contingency.

**Figure 4.**
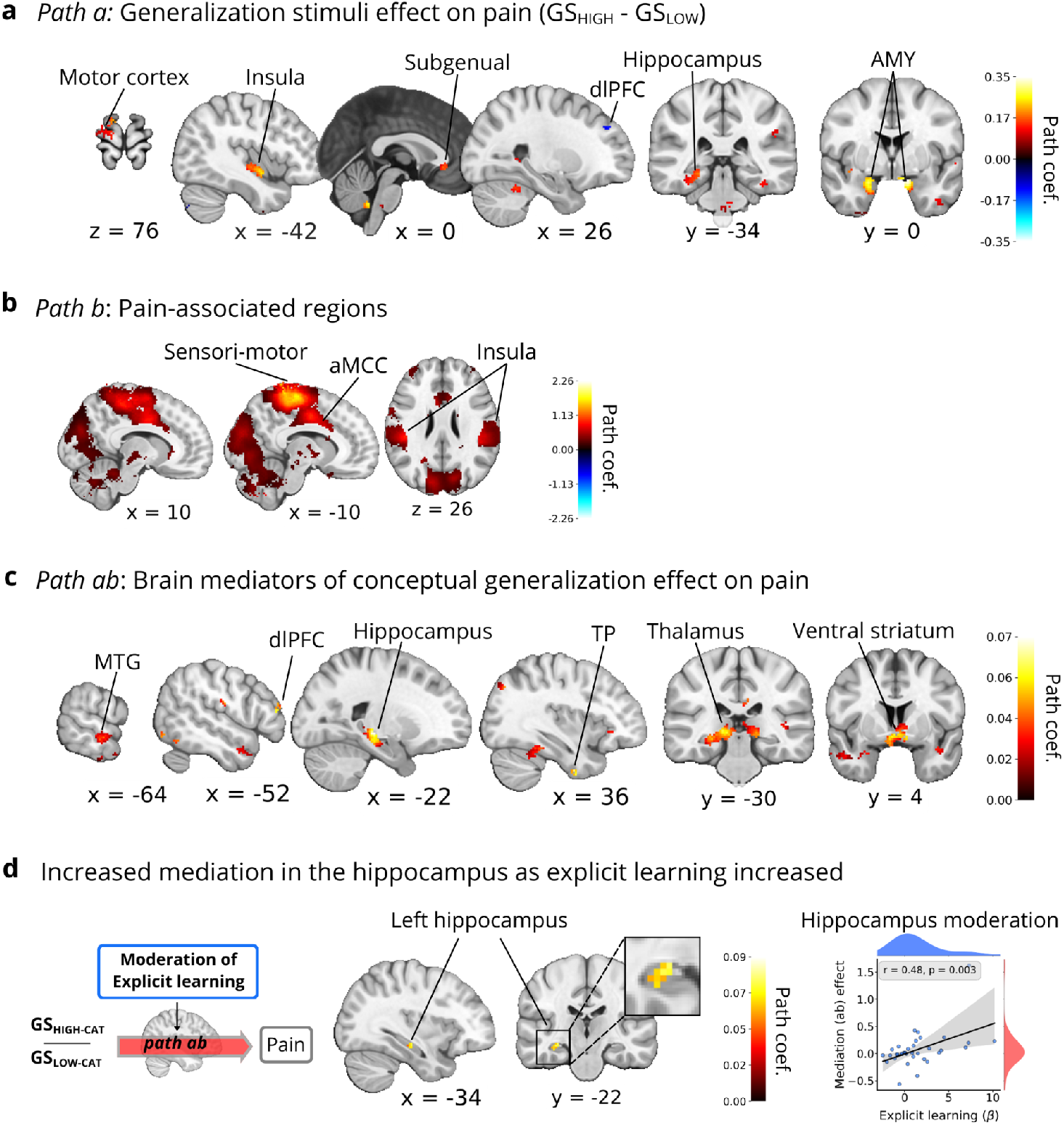
Multilevel mediation analysis of conceptual generalization of pain. All results are displayed at an FDR corrected threshold of *q* < 0.05. **a)** Effect of generalization condition on pain-related brain activity (*Path a*). Regions showing significantly greater activity during pain for GS_HIGH-CAT_ compared to GS_LOW-CAT_ trials included the bilateral amygdala, left mid-posterior insula, subgenual anterior cingulate cortex (sgACC), left posterior hippocampus, and right motor cortices. A small cluster in the right dorsolateral prefrontal cortex (dlPFC) showed the opposite pattern, with greater activity for GS_LOW-CAT_. **b)** Pain-associated brain activity (*Path b*). Regions where trial-by-trial activation during pain was positively associated with subjective pain ratings (controlling for generalization condition) included the bilateral anterior and posterior insula, anterior midcingulate cortex (aMCC), thalamus, primary and secondary somatosensory cortices, motor areas, brainstem, cerebellum, left dlPFC, bilateral visual areas and bilateral superior parietal cortex. **c)** Brain mediators of conceptual generalization effects on pain (*Path ab*). Regions where activation formally mediated the effect of generalization condition on pain ratings included bilateral ventral striatum (including nucleus accumbens and caudate), thalamus, left hippocampus, left temporopolar cortex, lateral and anterior frontal poles, supplementary motor cortex, bilateral mid-temporal gyri, and bilateral posterior parietal cortex. Activation in these regions for GS_HIGH-CAT_ > GS_LOW-CAT_ trials was associated with increased pain ratings. **d)** Moderation of hippocampal mediation by explicit learning. The strength of hippocampal mediation (*Path ab*) was positively correlated with individual differences in explicit learning of CS-pain associations. This suggests that hippocampal mediation of conceptual generalization effects on pain is stronger for individuals who have developed explicit expectations about the pain-predictive value of the CSs. The scatterplot illustrates this positive relationship for visualization purposes only. Each dot represents a subject.

*Path b* tested which brain regions are associated with trial-by-trial variations in subjective pain ratings, controlling for stimulus type (GS_HIGH-CAT_ > GS_LOW-CAT_). As expected, pain-related brain activity was observed in regions classically associated with pain processing (*q* < 0.05 FDR corrected, Figure 4b), including positive associations in the bilateral anterior and posterior insula, anterior midcingulate cortex (aMCC), thalamus, primary and secondary somatosensory cortices, motor areas, brainstem, and cerebellum. Additional positive effects were observed in the left dlPFC, bilateral visual cortices, and, to a lesser extent, the bilateral superior parietal cortices and left temporal pole. These findings confirm that increased activation in pain-related and higher-order regions was related to higher subjective pain ratings, independent of stimulus type.

*Path ab* identifies brain regions that formally mediated the relationship between generalization stimuli category and pain ratings. Significant positive mediation effects (*q* < 0.05 FDR corrected) were found across a distributed network, including areas associated with semantic and conceptual processing, memory and valuation. Strongest effects were found in the bilateral ventral striatum (including the nucleus accumbens and caudate), thalamus, left hippocampus and left temporopolar cortex. Additional effects were found in frontal regions, including the lateral and anterior frontal poles and supplementary motor area. Moreover, regions within the default mode network (Alves et al., 2019) also showed significant positive mediation effects, notably the left retrosplenial cortex, bilateral middle temporal gyri, and bilateral posterior parietal cortex. Finally, mediation effects were observed in the precuneus and cerebellum, particularly in crus I, crus II, and lobule IV.

In addition to the mediation effect observed in the left hippocampus, this area also showed a significant, positive moderation effect by explicit learning, as indicated by a positive correlation between the person-level *ab* effect and individual differences in pain expectation (CS_HIGH_ - CS_LOW_) during the associative learning task (Figure 4d). Thus, participants who had explicitly learned the prior CS-pain associations showed stronger hippocampal mediation effects of generalization cue category on pain.

### Representational similarity analysis

To further characterize the representational structure of hippocampal activity during conceptual pain generalization, we conducted an exploratory representational similarity analysis (RSA) in the left hippocampus. We focused on the left hippocampus as it was the only region in which pain-evoked activity during generalization mediated the effects on pain ratings and was moderated by explicit learning. We compared two representational models: an exemplar model, which predicted greater similarity between stimuli depicting the same exemplar across modalities, and a categorial model, which predicted greater similarity between stimuli belonging to the same pain-related category (GS_HIGH_ or GS_LOW_; Figure 5a).

**Figure 5.**
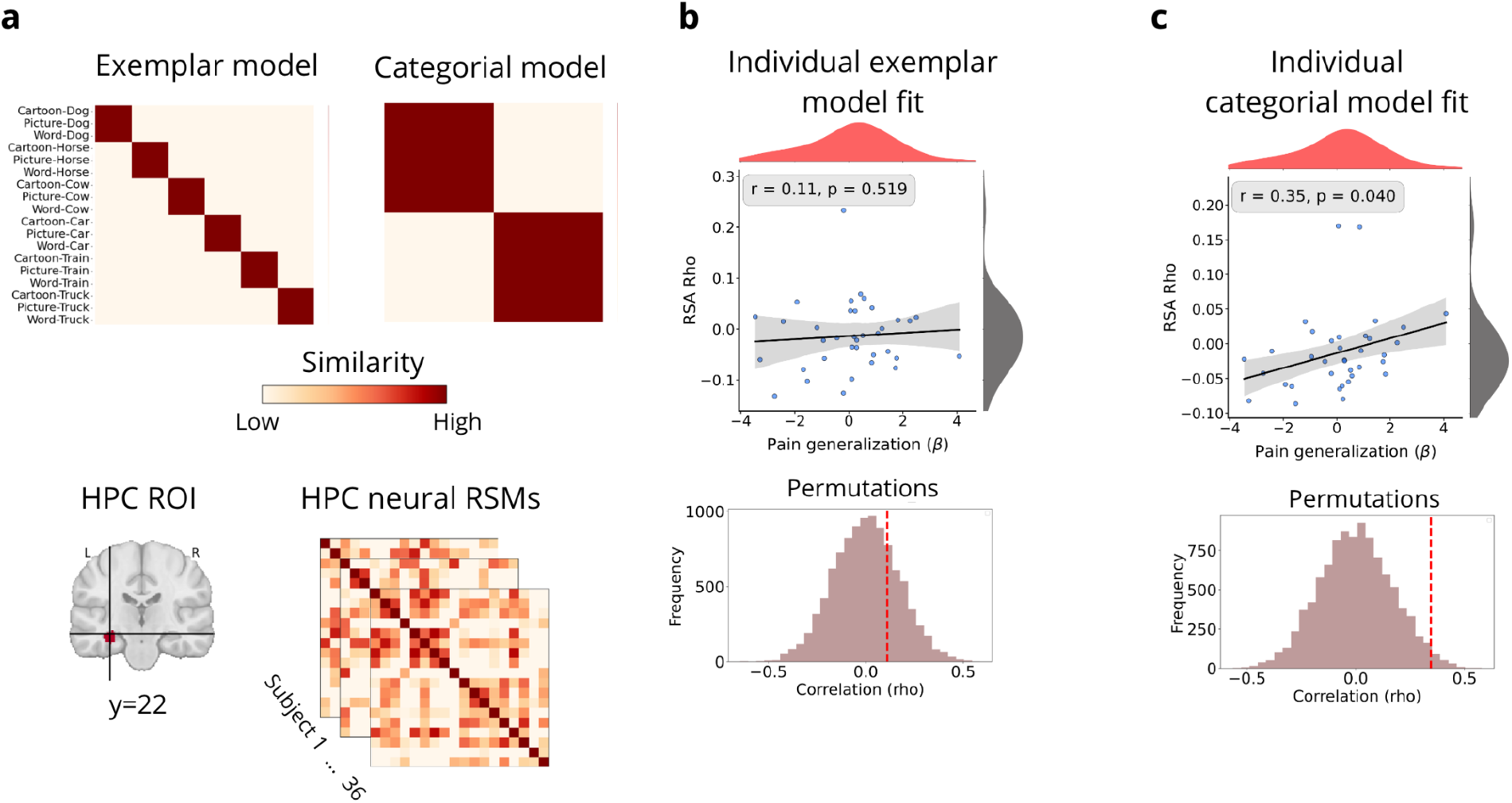
Representational similarity analysis in the left hippocampus. **a)** Representational models and hippocampal representational similarity matrices. The exemplar model assigns greater similarity between generalization stimuli (GS) depicting the same exemplar across different modalities (e.g., dog presented as a word, drawing, or image). The category model assigns similarity between GS belonging to the same pain-related conceptual category (GS_HIGH-CAT_ or GS_LOW-CAT_), regardless of the exemplar. Individual hippocampal representational similarity matrices were computed from pain-evoked activity within the left hippocampal ROI (MNI coordinates: −32, −22, −12). The fit between each model and individual hippocampal similarity matrices was assessed with Spearman rank correlation, and p-values were obtained by permuting the rows and columns of the neural matrix 10,000 times before recomputing the fit. The mean RSA fit at the group level was not greater than zero for both the exemplar and the categorial model. **b)** Association between individual RSA model fits and behavioral conceptual pain generalization effects. Correspondence with the exemplar model was not associated with behavioral generalization (*r* = .11, *p* = .52; 10,000 permutations). **(c)** Stronger behavioral generalization was associated with higher RSA values for the categorial model (*r* = .35, *p* = .040; 10,000 permutations). Scatterplots show individual participants. Histograms show the null distributions generated by permutation testing, with the observed correlation indicated by the red dashed line.

Neither the exemplar nor the categorial models described the hippocampal representational structure well at the group level (t(35) = -1.22, *p* = .23 and t(35) = -1.37, *p* = .18, respectively). However, individual differences in category-based representational structure were positively associated with individual differences in the behavioral effect of conceptual pain generalization, such that participants who showed greater correspondence between their hippocampal representational structure and the categorial model also showed greater behavioral generalization effects (*r* = .35, *p* = .04; 10,000 permutations; Figure 5c). In contrast, correspondence with the exemplar model was not significantly related to behavioral generalization (*r* = .11, *p* = .52; Figure 5b). Of note, negative categorial RSA fits were observed in several participants. A negative fit indicates a representational structure opposite to that predicted by the category-based model, with relatively greater similarity for between-category than within-category comparisons.

## Discussion

Pain is shaped by prior expectations and predictions, often formed through learned associations between events and painful experiences. These expectations can generalize beyond the original learning setting, biasing how pain is perceived and processed in the brain. Conceptual generalization—defined as the transfer of learned associations based on abstract or high-level similarity—offers a framework for understanding this process. Clarifying how conceptual generalization shapes pain, both behaviorally and neurally, is important, as it may contribute to maladaptive responses such as nocebo effects and to chronic pain (Kampermann et al., 2021; Meulders, 2019; Weng et al., 2025).

In this study, we examined whether novel stimuli that share a conceptual resemblance to pain-predictive cues, but were never paired with any level of pain, could modulate pain perception, and used fMRI to probe the brain mechanisms underlying this effect. Although these generalization stimuli (GS) did not affect self-reported pain ratings at the group level, participants who had explicitly learned the original stimulus-pain associations reported greater pain in response to GS from the same conceptual category as high-pain learning stimuli. These behavioral effects were mirrored in the brain. Pain-evoked activity in the hippocampus was higher for generalization stimuli of the same conceptual category as the high-pain learning cue and statistically mediated the conceptual generalization effect on pain ratings. This effect was stronger among participants who had developed more explicit expectations. Additional mediation effects were observed in brain areas related to valuation, such as the ventral striatum, and in areas related to semantic processing, especially regions of the default mode network. Next, we specifically discuss the role of the hippocampus, given the theory-driven predictions of its involvement in conceptual generalization and in abstract inferential processes, followed by a system-level integration with other mediators of conceptual generalization.

Our work builds on previous behavioral studies (Koban et al., 2018; Liu et al., 2019) showing that altered pain reports to novel stimuli are not automatic. Instead, they depend on contingency awareness, whether people have explicitly learned the initial stimulus-pain associations during conditioning. This suggests that conceptual generalization reflects higher-order inference rather than passive or implicit processes (Dunsmoor & Murphy, 2015; Kumaran, 2012). We further show that both behavioral and brain responses to generalization stimuli correlate with the strength of prior learning. Participants who formed stronger stimulus-pain expectations during conditioning tended to exibit greater generalization effects and stronger hippocampal modulation during pain.

Fear learning studies show that contingency awareness moderates conditioning (Baeuchl et al., 2019; Klucken et al., 2009; Mertens, Boddez, et al., 2021). Our findings extend this to conceptual generalization, suggesting that generalization across contexts is, for the most part, conditional on explicit knowledge of cue-outcome relationships. This mechanism aligns with research showing that pain-related expectations generalize across various settings (Glogan et al., 2023; Kloos et al., 2022; Mertens et al., 2021; Meulders et al., 2017; Meulders & Bennett, 2018). Our results support a cognitive mechanism grounded in abstract representations and explicit expectations influencing pain processing.

Brain mediation analyses identified the left mid-to-posterior hippocampus as a key region linking conceptual generalization to pain modulation. It was more activated during pain following a generalization stimulus conceptually related to CS_HIGH_ versus CS_LOW_ (*Path a*), and statistically mediated the effects of stimulus category on increased pain reports (*Path ab*), especially for participants who had developed explicit CS-pain expectations during learning (moderation of *Path ab* effect). While the anterior hippocampus encodes predictive value of pain-associated cues (Wimmer & Büchel, 2021) and contextual pain features (Caston et al., 2023), the mid-posterior segment supports associative processing, memory retrieval, and perceptual integration (Kim et al., 2015; Robinson et al., 2015). Our results align with models of the hippocampus as a hub for cognitive maps, in which abstract relational representations (e.g., category membership) support inference in novel situations (Constantinescu et al., 2016; Garvert et al., 2017; Sherrill et al., 2023). Garvert et al. (2017) implicated the mid-posterior hippocampus in such inferential processes, consistent with our peak mediation and moderation effects.

In predictive processing models of pain, prior expectations shape incoming nociceptive signals, amplifying or attenuating pain based on learned associations and expected value (Seymour, 2019). Our findings support the idea that the hippocampus may contribute to this predictive coding-based pain modulation, by linking cues to knowledge-based conceptual maps. Such processes could help explain how pain-related expectations persist, generalize, and influence new experiences. This mechanism has implications for chronic pain. The fear-avoidance model (Vlaeyen, 2015; Vlaeyen & Linton, 2000) describes how fear of pain leads to avoidance behavior, which can generalize to similar contexts or movements. This generalization of pain-related behavior and cognition can block extinction of maladaptive behaviors and recovery, ultimately contributing to the maintenance of pain (see Meulders, 2019, for a review). Our findings suggest that the hippocampus may support this process by enabling conceptual generalization of learned expectations.

Supporting this, animal models have identified the hippocampus as a critical site in the development of chronic pain. Blocking neurogenesis in this region prevents the transition to chronic pain after nerve injury (Apkarian et al., 2016; Mutso et al., 2012, 2014). In humans, altered hippocampal-prefrontal connectivity has been linked to pain chronification (Vachon-Presseau et al., 2016). More recently, the functional connectivity of the posterior hippocampus with other posterior regions predicted chronic pain development after an injury in individuals without a prior history of chronic pain, potentially reflecting altered affective learning and expectations in these individuals (Branco et al., 2024). Together, these findings suggest that the hippocampus may be a key site where pain regulation, learning processes and conceptual mechanisms interact.

Several features of our paradigm help constrain the interpretation of these findings. Generalization stimuli were never directly paired with pain and were presented in three distinct modalities (images, drawings, and written words), minimizing perceptual overlap (see Figure 1b). Their only shared feature was a conceptual category (e.g., animal or vehicle) with the initial pain-predictive conditioned stimuli. Brain activity was measured during pain *itself*, not during cue presentation or anticipation, ensuring that hippocampal activation reflected the abstract pain value inferred from conceptual similarity to prior learning. Thus, the observed hippocampal differences are consistent with the involvement of conceptual information during pain processing, rather than reflecting only anticipatory responses to the generalization cues.

Our exploratory RSA further suggests that, although non-significant at the group level, individual differences in hippocampal representational structure may be related to category-based conceptual generalization effects on pain. Participants who showed stronger behavioral generalization tended to show greater correspondence between hippocampal activity patterns and the categorical model, whereas participants showing weak or absent generalization tended to show the opposite representational pattern, with relatively greater dissimilarity among stimuli within the same pain-related category (i.e., GS_HIGH-CAT_ - GS_LOW-CAT_). This pattern raises the possibility that individual differences in how the hippocampus organizes or differentiates conceptually related pain representations may contribute to individual differences in the generalization of learned pain modulation. The greater categorial similarity observed in stronger generalizers and the greater within-category differentiation observed in weaker generalizers are broadly compatible with complementary processes analogous to pattern completion and pattern separation, respectively. Pattern completion and pattern separation in the hippocampus have been associated with the integration and differentiation, respectively, of representations sharing overlapping features (LaRocque et al., 2013; Rolls, 2016; Yassa & Stark, 2011). Future studies using designs specifically developed to distinguish representational integration from pattern separation will be needed to test whether these hippocampal processes contribute to individual differences in the generalization of learned pain expectations.

Beyond the hippocampus, our mediation analysis identified a broader network involved in conceptual generalization, including the ventral striatum, thalamus, temporal pole, and default mode network (DMN). DMN regions, involved in semantic integration and self-referential processing, likely support top-down inferences linking conceptual knowledge to sensory input (Koban et al., 2021; Menon, 2023). Their recruitment aligns with evidence that the DMN enables context-specific decision-making and inference based on learned rules (Bowman & Zeithamova, 2018; Vatansever et al., 2017; Zeithamova & Bowman, 2020). DMN-hippocampal interactions also support memory retrieval and recollection of prior experiences (Raichle, 2015; Spreng et al., 2010).

The ventral striatum also emerged as a significant mediator, suggesting that valuation and motivational systems contribute to conceptual generalization of pain modulation. While typically linked to reward learning, the striatum has also been implicated in aversive learning (Klein et al., 2022) and in pain valuation (Baliki et al., 2010, 2013). Mediation in the hippocampus and striatum partially overlapped with ‘pro-pain’ regions in the SIIPS1 model of endogenous pain modulation, but also showed overlap with ‘anti-pain’ regions (Woo et al., 2017). Hippocampal-striatal interactions support generalization and inference in reward learning (Colas et al., 2022; Garvert et al., 2023; Wimmer & Shohamy, 2012). Our results suggest that conceptual representations in the hippocampus interact with affective value via striatal circuits during generalization. These results also raise the possibility that hippocampal-striatal interactions in pain generalization reflect the influence of explicit expectations, consistent with previous findings of striatal modulation by explicit conceptual influences (such as instructions) in reinforcement learning and decision-making (Koban et al., 2017; Li et al., 2011; Staudinger & Büchel, 2013).

In contrast, regions involved in threat detection and somatosensory processing, such as the amygdala, mid-posterior insula, and motor cortex, showed increased activation to generalization stimuli conceptually related to CS_HIGH_ compared to CS_LOW_. However, these regions did not mediate the effect of stimulus category on pain ratings or interact with individual differences in explicit learning. Their activity likely reflects generalized threat or salience responses rather than expectation-based modulation, consistent with findings that the amygdala responds more to conditioned than to instructed cues (Atlas et al., 2016; Koban et al., 2019), and supports perceptual threat generalization (Lissek et al., 2014; Webler et al., 2021).

Together, these findings point to a dual-mechanism account of conceptual generalization in pain. An aversive-sensory circuit (amygdala, insula, motor cortex) encodes generalized high-pain representations that may remain unconscious, while in parallel, a higher-order conceptual network (hippocampus, striatum and DMN) retrieves abstract category-based expectations that may shape pain through top-down modulation.

Our findings suggest a novel mechanism by which nocebo effects, i.e., increased pain driven by expectations, may generalize to new cued contexts. Previous studies have shown that the hippocampus interacts with pain-related networks when more pain is expected, and that this activity correlates with individual differences in pain expectations (Bingel et al., 2011, 2022; Jensen et al., 2015; Kong et al., 2008). These effects are typically observed in response to direct conditioning, and they tend to generalize most strongly across perceptually similar cues or environments (Kampermann et al., 2021; Weng et al., 2025). We extend this by showing that pain responses generalize across conceptually related, but perceptually distinct stimuli, even without direct pain pairing. Our results mirror nocebo mechanisms, where expectations amplify pain without changes in nociceptive input. This hippocampal-driven generalization may explain why people experience pain in novel settings symbolically linked to prior pain.

In addition to hippocampal contributions, other regions involved in conceptual processes, self-referential and social cognition, particularly within the DMN, may support other forms of expectancy-based pain modulation. DMN regions have been implicated in integrating socially and conceptually acquired pain-related expectations, such as through observational learning (Schenk & Colloca, 2020) and interpersonal interactions (Ellingsen et al., 2023). Our findings may inform broader research on how high-level influences shape pain experience.

Future work could examine conceptual generalization in chronic pain populations, where greater generalization of pain and pain-related fear is expected (Meulders et al., 2014, 2017). We expect stronger DMN mediation in these individuals, consistent with evidence of heightened self-referential processing (Fiúza-Fernandes et al., 2025). Also, we used expectation ratings during learning but not during generalization. Future studies could include pain ratings to better assess expectations and allow computational modelling of conceptual generalization. We also note that the association between learning and generalization does not by itself establish why some individuals learn less during the initial learning phase. Thus, weaker learning and subsequent generalization of cue-pain associations could reflect differences in attention, sensory discrimination of heat pain stimuli, or discrimination of both learning and the generalization cues, rather than differences in the transfer of learned pain expectations per se. The present design cannot fully distinguish these possibilities. While the empirical similarity structure suggests that the generalization stimuli and categories were discriminable, future studies could independently assess stimuli discrimination during learning and category recognition during generalization. This would help to determine whether individual differences in pain generalization primarily reflect differences in learned expectations and their transfer to related conceptual cues, differences in category-level representation, or interactions between these processes. Finally, we note that the generalization categories in our study were restricted to animal and vehicle categories. Testing broader and more ecologically valid categories, such as real-life actions (Glogan et al., 2023) or social cues (Koban et al., 2019), would be beneficial to clarify how conceptual generalization shapes pain in everyday contexts.

In conclusion, this study shows that stimulus-pain associations can generalize based on abstract conceptual relationships and bias how pain is experienced. Novel stimuli that were conceptually, but not perceptually, related to pain-predictive cues increased pain ratings, especially in participants who had developed more explicit expectations during initial learning. This behavioral effect was mirrored in the hippocampus, where pain-evoked activity reflected conceptual generalization, predicted pain ratings, and scaled with expectation strength. A broader network related to valuation and conceptual processing also contributed, while threat-related regions responded to conceptual generalization stimuli but were not linked to subjective pain ratings. Together, these findings highlight how conceptual associations can shape pain in novel contexts through expectations, with implications for understanding the generalization of nocebo effects and maladaptive pain responses.

## Methods

### Participants

A total of 38 healthy volunteers, with no previous psychiatric, neurological, or pain disorders, took part in the study. Two participants were excluded from the analyses due to high movement artifacts or technical complications during the experiment. The final sample comprised 36 individuals (20 females and 16 males, mean age = 27.1 years, age range 18–50 years). All participants provided informed consent and received compensation for their time. The study adhered to the Declaration of Helsinki and was approved by the Institutional Review Board at the Department of Psychology and Neuroscience, University of Colorado Boulder.

The design of this fMRI study was informed by three previously reported behavioral experiments demonstrating a consistent moderating role of individual differences in learning-related expectations on pain generalization (Koban et al., 2018). Across these studies (total N = 134), which included both perceptual and conceptual generalization paradigms, the corresponding interaction effects between learning and generalization were of moderate effect sizes. Based on the interaction statistics reported in these studies, we estimated a pooled effect of approximately *r* = .54. A two-tailed power analysis (α = .05, 90% power) indicated that approximately 32 participants would be required to detect an effect of this magnitude.

### Stimuli

The learning stimuli (conditioned stimuli, or CSs) consisted of three cartoon images of animals and three of vehicles. For each participant, only one animal and one vehicle image were shown during the learning phase, with one of them assigned to the high-pain condition (CS_HIGH_) and the other to the low-pain condition (CS_LOW_). The two learning stimuli and the assignment to high versus low pain were completely counterbalanced across participants. The generalization stimuli (GS) comprised 18 novel cues that were not shown during the learning phase. These included three animals (cow, horse, dog) and three vehicles (car, train, truck). Each concept was represented in three different visual display modalities (drawings, photographs, and written word), resulting in nine GSs per category. GSs were categorized based on their conceptual relationship to the CS assigned to high or low pain. The set of GSs conceptually related to the category of the CS_HIGH_ was labeled GS_HIGH-CAT_, and the ones conceptually related to the category of the CS_LOW_ were labeled GS_LOW-CAT_.

### Materials and procedure

Participants first completed a classical conditioning task designed to induce pain modulation through learned associations by pairing visual stimuli (cartoon images of animals or vehicles) with different levels of noxious heat (see Figure 1a: *Associative Learning*). One stimulus (CS_LOW_) was paired with low to medium heat (48°C or 49°C), and the other (CS_HIGH_) with medium to high heat (49°C or 50°C). Conditions (CS_HIGH_ versus CS_LOW_, e.g., animal versus vehicle) were counterbalanced across participants and selected from sets of three cartoons to reduce image-specific effects (see *stimuli*). To isolate the effect of learned pain modulation, medium-intensity heat (49°C) was used on half the trials with each CS. This design allowed comparison of pain responses to identical medium-intensity heat stimulation as a function of CSs. On each trial, participants first rated their expected pain on a visual analog scale (VAS; 0-100), anchored at “no pain at all” to “worst pain”. After receiving the heat stimulus, they rated their experience of pain using the same scale. The learning task included 96 trials (for details, see Koban et al., 2019).

In the generalization phase, participants were shown novel visual stimuli that were conceptually related to the CSs but had not been shown previously. Half of the 18 GS were conceptually related to each participant’s CS_HIGH_, and half to their CS_LOW_ (Figure 1a: *Conceptual generalization task*). Each GS was presented for 3 seconds, followed by a jittered interval (2–4s). They were then followed by a medium-intensity heat stimulus (49°C), delivered on the right leg, using a 27mm diameter fMRI-compatible CHEPS thermode monitored by a Medoc Pathway system (Medoc, Israel). For each trial, after the heat stimulus, participants rated their pain using the same VAS as in the learning phase. The generalization task included 72 trials, divided into four scanning runs of 18 trials each. The thermode location was changed between each run to prevent sensitization.

At the end of the experiment, participants completed a similarity rating task involving all visual stimuli used in the experiment, including CSs that were not part of their own learning phase (see *stimuli*). They were asked to rate how similar each GS was to each CS, using a VAS ranging from “not similar at all” to “completely similar”. No mention was made of visual versus conceptual similarity in the instructions to avoid biasing the assessment. These ratings were used to validate that GSs and CSs from the same conceptual category (e.g., animals or vehicles) were perceived as similar (see *conceptual similarity* below). Due to a technical error, eleven participants only rated half of the potential combinations between GSs and CSs. Therefore, the group average similarity matrix was used for further analyses.

### fMRI acquisition and preprocessing

Functional brain activity was recorded using a Siemens TrioTim 3T scanner, capturing the brain in 26 interleaved transverse slices with 3.4mm isotropic voxels. Images were acquired with a T2* weighted EPI GRAPPA sequence (TR = 1.3 s, TE = 25ms, flip angle = 50°, FOV = 220 mm) and preprocessed using SPM, following a standard procedure that included motion correction, slice-timing correction, spatial normalization to MNI space, and spatial smoothing with an 8-mm FWHM Gaussian kernel. Preprocessed images were resampled to 3×3×3 mm voxel size. For spatial normalization, T1-structural MPRAGE images (1 mm isotropic voxels) were first coregistered to the mean functional image and then normalized to the SPM template using unified segmentation.

### Statistical analyses

We used multilevel general linear models (GLMs) to examine how experimental conditions influenced pain ratings. All models included participant-specific random intercepts and slopes to account for individual differences in baseline pain sensitivity and in learning rates. Code for all analyses is available at https://github.com/canlab.

### Learning task

To test whether the learning task resulted in modulation of pain, we focused on trials with medium-intensity heat stimulation. This design allowed a direct comparison of pain ratings for identical stimuli as a function of stimulus type (CS_HIGH_ versus CS_LOW_, coded as 1 and -1). A significant difference between CS_HIGH_ versus CS_LOW_ trials indicated that pain ratings were influenced by the learned pain predictive value of each cue.

We also tested whether stimulus type (CS_HIGH_ versus CS_LOW_) influenced pain expectation ratings for medium-intensity trials. The multilevel GLM estimated a random slope for each participant, reflecting how strongly expected pain was influenced by the stimulus type. These participant-level slopes (beta estimates) were used as a between-person measure of explicit learning and were used as moderators for the analysis of behavior and fMRI data in the subsequent generalization task, in line with the results from a series of three previous behavioral studies (Koban et al., 2018) that have shown that explicit learning plays a key role in the generalization of pain modulation.

### Generalization task

The goal of the conceptual generalization task was to test whether pain ratings were modulated by the category of the generalization stimulus, i.e., whether cues conceptually related to CS_HIGH_ or CS_LOW_ modulated pain responses to identical noxious stimuli. We used multilevel GLMs to estimate the effect of the generalization stimulus category (GS_HIGH-CAT_, GS_LOW-CAT_, coded as 1 and –1) on pain ratings. Explicit learning scores from the associative task were included as a second-level moderator to test whether individual learning differences modulated the generalization effect. Because this moderation effect constitutes the primary behavioral finding motivating the subsequent neural analyses, we additionally assessed its robustness using a non-parametric permutation procedure. Specifically, we randomly permuted the participant-level explicit learning scores across participants 5,000 times and recomputed the multilevel GLM and corresponding moderation statistic for each permutation. The resulting null distribution was used to evaluate the significance of the observed moderation effect without relying only on the assumptions of the parametric model.

### Conceptual similarity

To characterize the structure of perceived similarity between generalization and conditioned stimuli, we analyzed participants’ similarity ratings collected at the end of the experiment (see *Materials and procedure*). Participants rated how similar each GS was to each CS, including CSs not seen during their own learning phase. No reference was made to conceptual characteristics or visual features in the instructions.

To reduce the influence of missing data caused by a software malfunction, we computed an empirical similarity matrix using the median rating, instead of the mean, across participants for each GS-CS pair (see Figure 1b). Visual inspection of this matrix suggested that participants grouped stimuli based on conceptual categories, and not modality. For example, the concepts of “dog,” “horse,” and “cow,” regardless of modality (word, picture, or cartoon), were rated as highly similar to animal CSs, but dissimilar to vehicle stimuli.

To formalize this observation and to characterize the structure of these empirical ratings, we compared the similarity matrix to three models using Spearman correlations (see Figure 1b). The exemplar model was a binary matrix assuming maximal similarity (1) only between CS-GS pairs that referred to the same object (e.g., dog word, dog picture, dog cartoon), and no similarity (0) to all other items, even if they belonged to the same conceptual category (e.g., animal). The category model also used binary coding but assumed maximal similarity between all items from the same conceptual category (e.g., all animals), regardless of modality or exemplar. Finally, the ResNet-50 visual model estimated similarity based on low-level image features, using cosine similarity between image embeddings of each CS and GS. ResNet-50 is a deep convolutional neural network designed for image classification and feature extraction (He et al., 2016; https://huggingface.co/microsoft/resnet-50). This network was chosen over other models because its architecture has been shown to simulate early and mid-level visual processing in the human brain (Schrimpf et al., 2018).

Among the three, the category model best explained the empirical ratings (*r* = 0.83, *p* < 0.001), outperforming both the exemplar model (*r* = 0.63, *p* < 0.001) and the ResNet-50 model (*r* = 0.21, *p* < 0.001). These results confirm that participants represented stimuli based on conceptual category rather than perceptual similarity or only based on object/examplar specificity, supporting our use of the GS_HIGH-CAT_ and GS_LOW-CAT_ categorization in both behavioral and brain analyses.

### fMRI analyses

We used SPM12 to estimate trial-by-trial pain-evoked brain activity using a single-event general linear model (GLM). Six motion parameters were included as covariates. This approach produced one beta map for each of the 72 generalization trials. Although single-trial models provide greater sensitivity to trial-level variation than block designs, they are more susceptible to noise. To address this, we calculated the variance inflation factor (VIF) for each trial to assess multicollinearity with regressors. Trials with a VIF greater than 4 were excluded, resulting in the removal of 33 beta maps (out of 2592 total). The remaining beta maps served as input for the multilevel mediation analyses. As in the behavioral analyses, we implemented multilevel models, estimating the effects of interest at the subject-level, before computing the group estimates, ensuring that the influence of noisy trials would be contained at the first level of the analysis.

We applied multilevel mediation effect parametric mapping using the CANlab Mediation Toolbox (https://github.com/canlab/MediationToolbox) to identify brain regions linking generalization conditions (GS_LOW-CAT_ and GS_HIGH-CAT_) to changes in pain rating. This approach models participants as random effects and tests whether a mediator (M) statistically explains the relationship between an independent variable (X) and an outcome (Y) (Atlas et al., 2010; MacKinnon et al., 2007). In our model, the GS category (coded as -1 for GS_LOW-CAT_ and 1 for GS_HIGH-CAT_) served as the independent variable (X), pain ratings as the outcome (Y), and pain-evoked beta maps as the mediator (M). As in behavioral analyses, explicit learning scores were included as a second-level moderator to test if mediation was influenced by individual differences in expectation during learning (see Figure 3).

The mediation model estimates three paths, each with a distinct interpretation. *Path a* tests whether the GS category influences brain activity. This path identifies voxels with significant differences in activation between generalization trials (GS_HIGH-CAT_ > GS_LOW-CAT_) and corresponds to a standard GLM contrast and reflects pain-related responses modulated by conceptual generalization. Positive or negative coefficients indicate whether a given region shows greater or lesser activation during GS_HIGH-CAT_ trials. *Path b* tests whether trial-by-trial variation in brain activity is associated with subjective pain ratings, controlling for GS category. This identifies pain-associated regions, regardless of the generalization condition. *Path ab,* the mediation path, tests whether changes in activity in a given voxel account for the effect of GS category on pain ratings. This identifies brain regions where activation during GS_HIGH-CAT_ versus GS_LOW-CAT_ trials explains increased pain ratings. Because this is a multilevel model, it captures both within- and between-subject effects. For example, positive mediation coefficients indicate that participants who show stronger activation for GS_HIGH-CAT_ also tend to report higher pain ratings on those trials.

We included participants’ explicit learning scores as a second-level moderator in the mediation model. The moderation tested whether individual differences in learning strength influenced the magnitude of mediation effects across the brain. We examined positive moderation effects for *Path ab*, identifying regions where the mediation was stronger among participants who developed explicit expectations about the pain-predictive values of CSs during the learning phase (see *Learning task*).

Significant clusters in each path were identified using False Discovery Rate (FDR) correction at *q* < 0.05. Across all analyses, the voxel-wise mediation models tested mediation effects in 586,836 voxels. For this number of comparisons, an FDR of *q* < 0.05 corresponded to *p* < 0.003. We report all significant clusters with a minimum size of *k* > 3 voxels.

We conducted an exploratory representational similarity analysis (RSA) to further characterize the representational structure of hippocampal activity during conceptual pain generalization. We focused on the left hippocampus because pain-evoked activity in this region was moderated by individual differences in explicit learning, paralleling the moderation observed behaviorally and identifying this region as the primary region showing an influence of learning on the generalization effect.

The RSA tested whether hippocampal activity patterns were organized according to individual stimulus exemplar or broader conceptual category structure. We constructed two a priori representational similarity models. The exemplar model predicted greater neural similarity between GS depicting the same exemplar across different modalities (e.g., the same dog presented as a word, drawing, or image). The category model predicted greater similarity between generalization stimuli belonging to the same pain-related conceptual category (GS_HIGH-CAT_ or GS_LOW-CAT_), regardless of the exemplar. For each participant, we extracted the mean multivoxel activation pattern for each of the 18 generalization stimuli by averaging across the four repetitions of each stimulus, yielding 18 stimulus-specific activation patterns per participant. These patterns were extracted from a 15-mm-radius sphere centred on the peak coordinate of the hippocampal mediation effect (MNI: −32, −22, −12) and anatomically constrained to the hippocampus according to the Julich-Brain atlas (Amunts et al., 2020), resulting in 235 voxels (Figure 5a). We then computed Pearson correlations between the activation patterns for every pair of generalization stimuli to generate an individual neural representational similarity matrix. For each participant, we quantified the correspondence between the neural representational similarity matrix and each model matrix using Spearman rank correlation of the upper triangle of each matrix. To assess the statistical significance of individual model fits, we performed 10,000 permutation tests for each participant and model by randomly permuting the rows and columns of the neural representational similarity matrix before recomputing the correlation with the model matrix. We assessed the mean fit of each model at the group level using a one-sample t-test against zero.

Finally, we tested whether individual differences in hippocampal representational structure were related to individual differences in behavioral conceptual pain generalization. Specifically, using Spearman rank correlation, we correlated each participant’s RSA model-fit estimate with their individual behavioral generalization effect, estimated as the within-participant coefficient for the generalization-category × explicit-learning interaction from the multilevel behavioral GLM (see *Behavioral analyses*). The significance of these correlations was assessed using 10,000 permutations.

## Contributions

L.K. and T.D.W. conceptualized the experiment. L.K. acquired the data. D.S.G. and L.K. analyzed the data. T.D.W. and L.K. obtained funding and supervised the project. D.S.G. and L.K. wrote the original draft and created the figures. All authors contributed to the interpretation of the data and to the editing of the paper.

## Conflicts of interest

The authors declare no conflict of interest.

## Acknowledgements

This study was funded by the National Institutes of Health (NIH; R37MH076136, PI: TDW) and an ERC Starting Grant SOCIALCRAVING (101041087, PI: LK). DSG was supported by Unifying Neuroscience and Artificial Intelligence and by Fonds de recherche du Québec—Nature et technologies excellence grants, as well as by Mitacs Globalinks fellowship programs. Views and opinions expressed are, however, those of the authors only and do not necessarily reflect those of the European Union or the European Research Council. Neither the European Union nor the granting authority can be held responsible for them. The funders had no role in study design, data analysis, manuscript preparation, or publication decisions.

## Supplementary information

**Supplementary Table 1.**
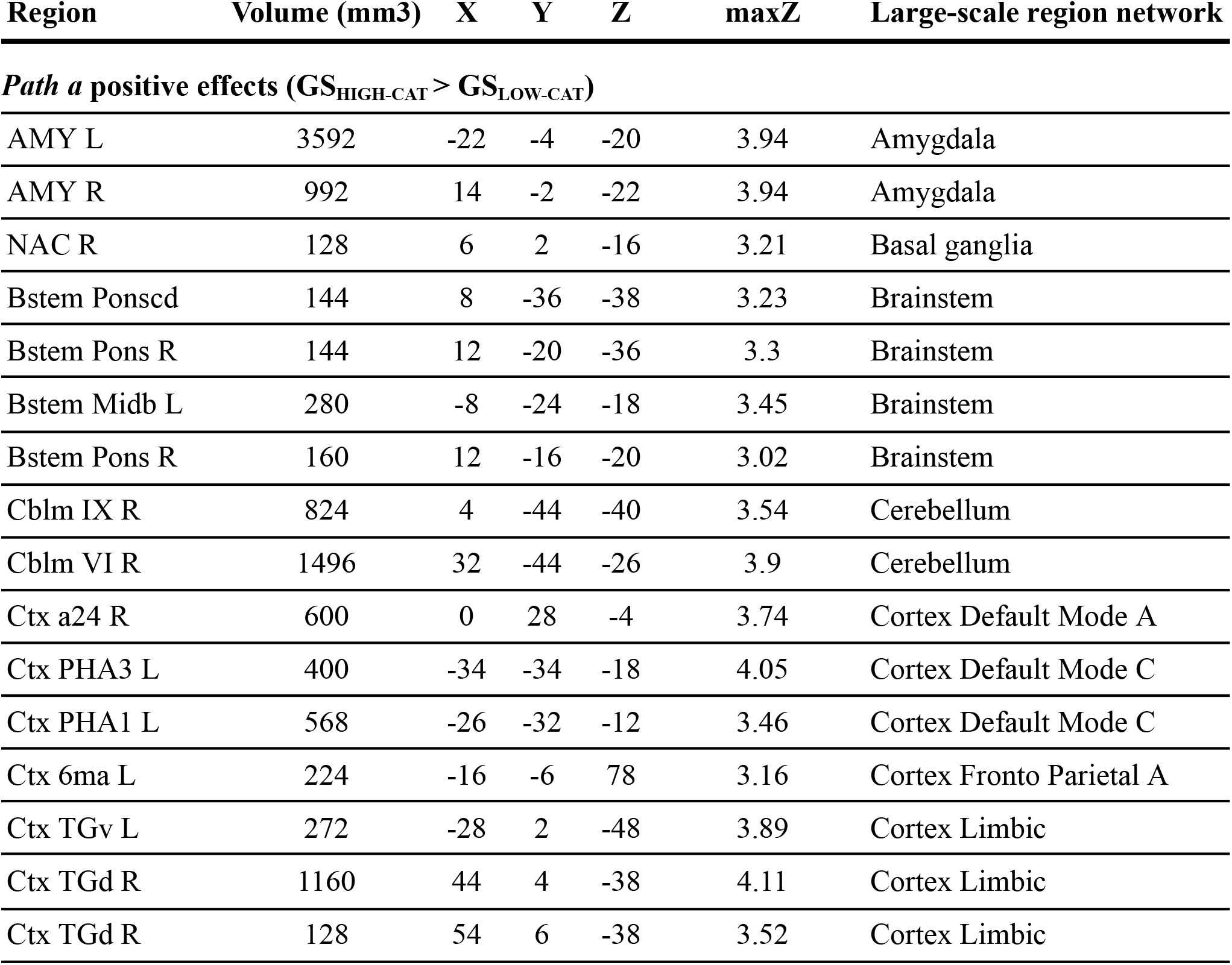
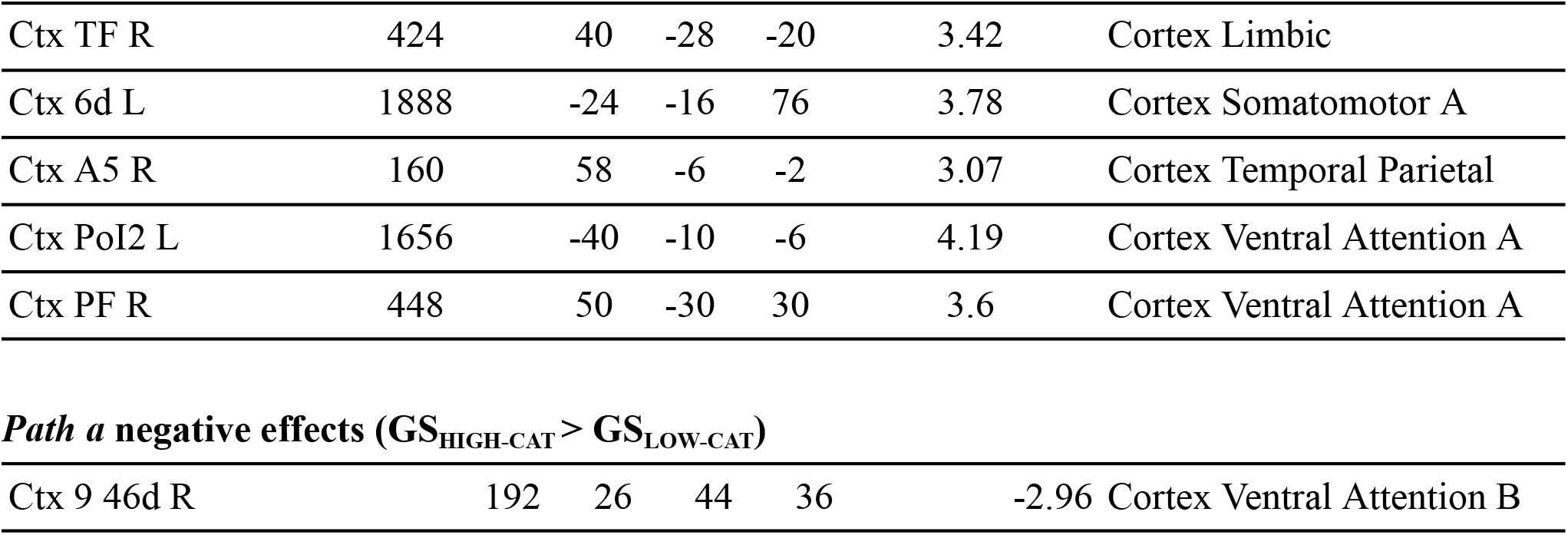
Path a effects from mediation analyses (FDR corrected, q < 0.05). This table reports positive and negative effects of the GS_HIGH-CAT_ > GS_LOW-CAT_ contrast during medium-intensity heat stimulation. Brain regions are labeled using the combined anatomical atlas developed by the CANlab group, which integrates several parcellation schemes. Cortical regions (Ctx) are based on 468 parcels from the multimodal Glasser et al. (2016) atlas, basal ganglia subdivisions from Pauli et al. (2016, 2018), cerebellar regions from Diedrichsen et al. (2009), brainstem subdivisions from Fairbust et al. (2007) and from Shen et al. (2013). The full combined anatomical atlas can be downloaded from https://github.com/canlab/Neuroimaging_Pattern_Masks/tree/master/Atlases_and_parcellations/2018_Wager_combined_atlas, with additional documentation available at https://sites.google.com/dartmouth.edu/canlab-brainpatterns/brain-atlases-and-parcellations/2018-combined-atlas. In addition, large-scale functional networks are labeled based on the atlas by Schaefer et al. (2018).

| Region | Volume (mm3) | X | Y | Z | maxZ | Large-scale region network |
| --- | --- | --- | --- | --- | --- | --- |
| ***Path a positive effects ($GS_{HIGH-CAT} > GS_{LOW-CAT}$)*** |  |  |  |  |  |  |
| AMY L | 3592 | -22 | -4 | -20 | 3.94 | Amygdala |
| AMY R | 992 | 14 | -2 | -22 | 3.94 | Amygdala |
| NAC R | 128 | 6 | 2 | -16 | 3.21 | Basal ganglia |
| Bstem Ponscd | 144 | 8 | -36 | -38 | 3.23 | Brainstem |
| Bstem Pons R | 144 | 12 | -20 | -36 | 3.3 | Brainstem |
| Bstem Midb L | 280 | -8 | -24 | -18 | 3.45 | Brainstem |
| Bstem Pons R | 160 | 12 | -16 | -20 | 3.02 | Brainstem |
| Cblm IX R | 824 | 4 | -44 | -40 | 3.54 | Cerebellum |
| Cblm VI R | 1496 | 32 | -44 | -26 | 3.9 | Cerebellum |
| Ctx a24 R | 600 | 0 | 28 | -4 | 3.74 | Cortex Default Mode A |
| Ctx PHA3 L | 400 | -34 | -34 | -18 | 4.05 | Cortex Default Mode C |
| Ctx PHA1 L | 568 | -26 | -32 | -12 | 3.46 | Cortex Default Mode C |
| Ctx 6ma L | 224 | -16 | -6 | 78 | 3.16 | Cortex Fronto Parietal A |
| Ctx TGv L | 272 | -28 | 2 | -48 | 3.89 | Cortex Limbic |
| Ctx TGd R | 1160 | 44 | 4 | -38 | 4.11 | Cortex Limbic |
| Ctx TGd R | 128 | 54 | 6 | -38 | 3.52 | Cortex Limbic |
| Ctx TF R | 424 | 40 | -28 | -20 | 3.42 | Cortex Limbic |
| Ctx 6d L | 1888 | -24 | -16 | 76 | 3.78 | Cortex Somatomotor A |
| Ctx A5 R | 160 | 58 | -6 | -2 | 3.07 | Cortex Temporal Parietal |
| Ctx PoI2 L | 1656 | -40 | -10 | -6 | 4.19 | Cortex Ventral Attention A |
| Ctx PF R | 448 | 50 | -30 | 30 | 3.6 | Cortex Ventral Attention A |
| ***Path a negative effects ( $GS_{\text{HIGH-CAT}} > GS_{\text{LOW-CAT}}$ )*** |  |  |  |  |  |  |
| Ctx 9 46d R | 192 | 26 | 44 | 36 | -2.96 | Cortex Ventral Attention B |

**Supplementary Table 2.**
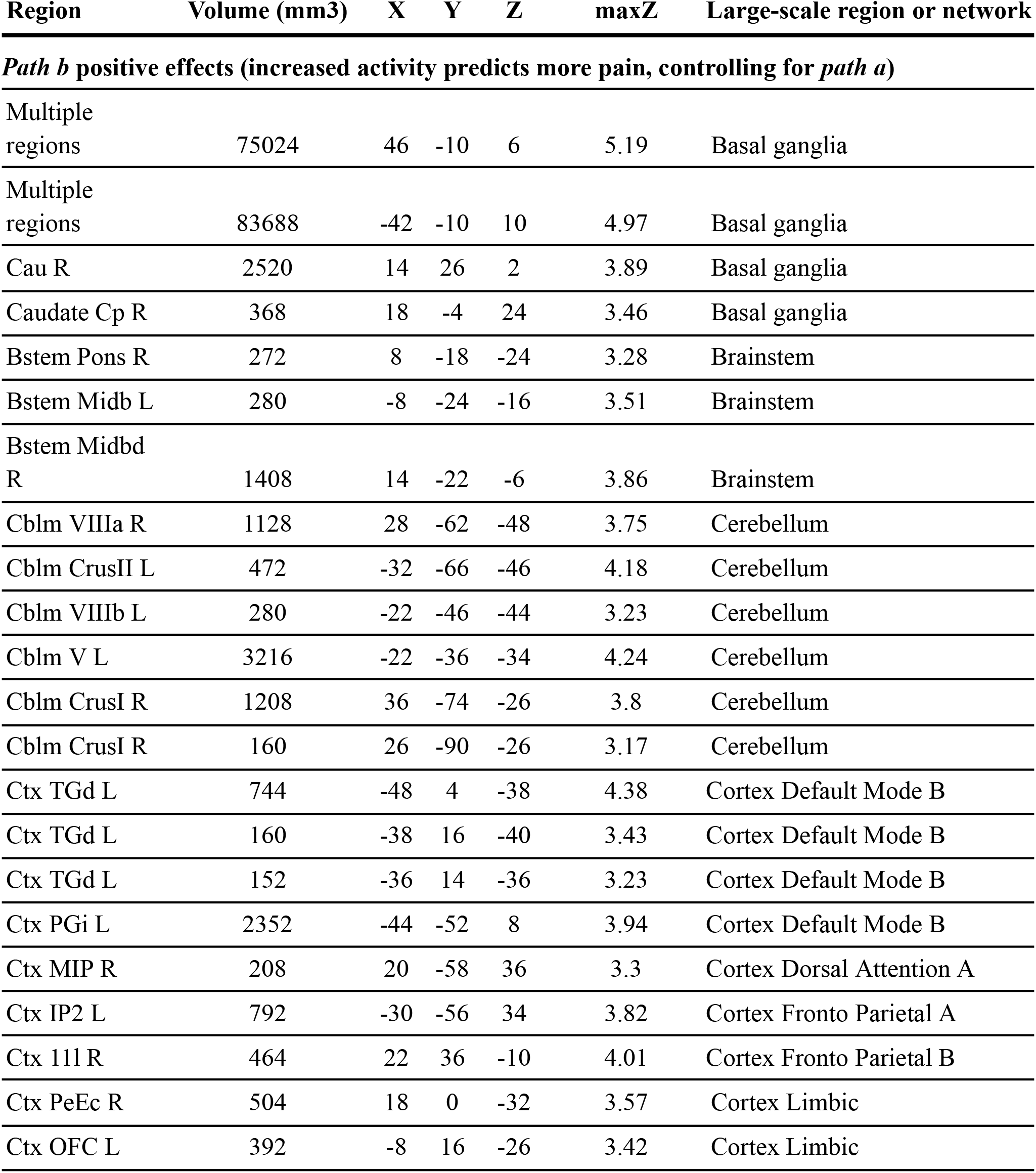
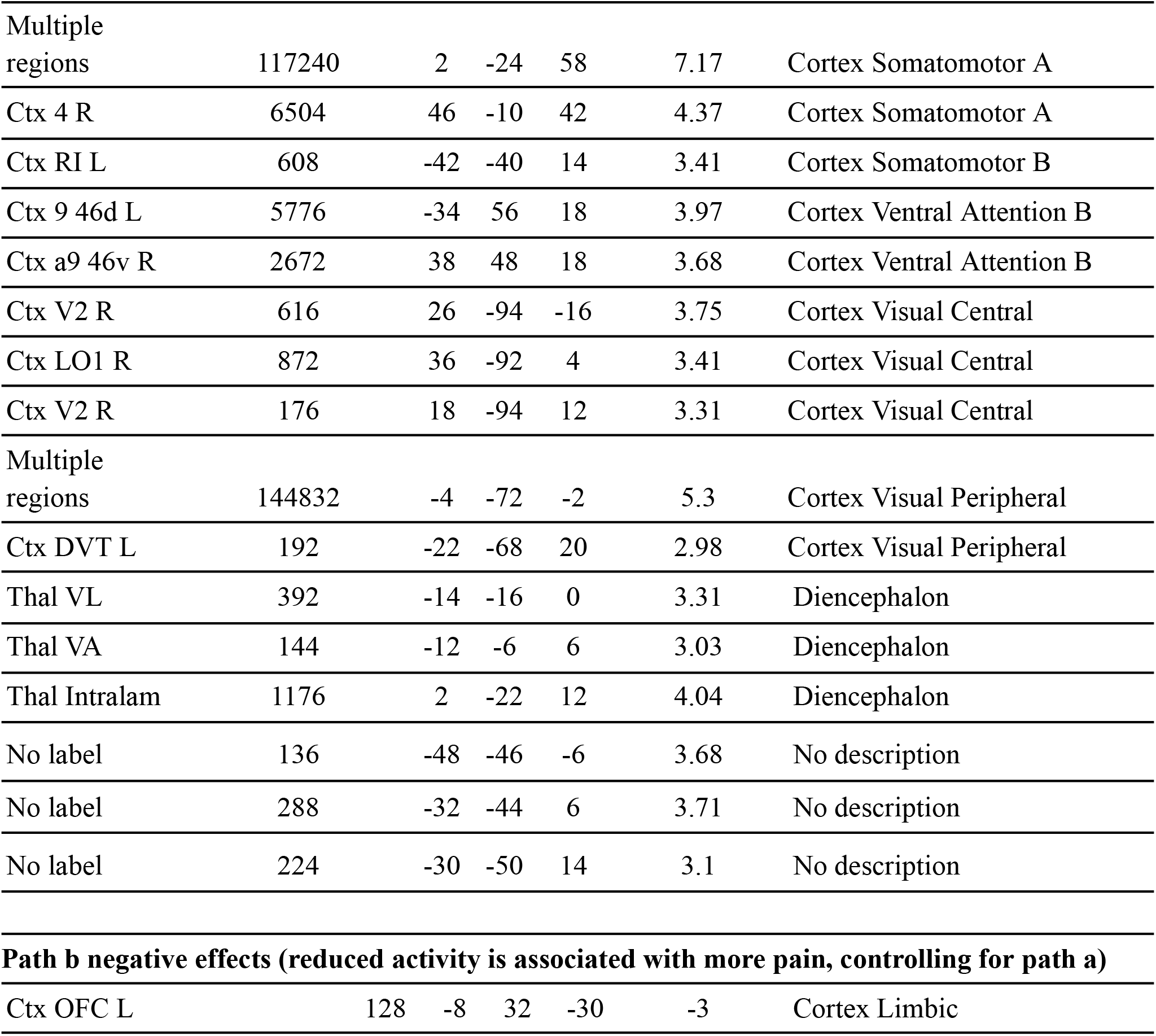
Path b effects of mediation analyses (FDR corrected *q* < 0.05). Positive and negative effects of brain activity associated with pain ratings on a trial-by-trial basis, while controlling for generalization conditions (GS_HIGH-CAT_ and GS_LOW-CAT_).

**Supplementary Table 3.** *Path ab* effects of mediation analyses (FDR corrected *q* < 0.05). Positive mediation effects indicate that increased pain-evoked activity during generalization (GS_HIGH-CAT_ > GS_LOW-CAT_) was associated with more pain ratings.

| Region | Volume (mm3) | X | Y | Z | maxZ | Large-scale region or network |
| --- | --- | --- | --- | --- | --- | --- |
| ***Path ab* positive effects (mediation of conceptual generalization effect on pain ratings)** |  |  |  |  |  |  |
| Multiple regions | 9376 | 2 | 4 | -6 | 5.64 | Basal ganglia |
| V Striatum R | 624 | 16 | 22 | -4 | 4.71 | Basal ganglia |
| Cau R | 616 | 24 | -32 | 14 | 5.03 | Basal ganglia |
| Caudate Ca R | 160 | 20 | 20 | 6 | 3.91 | Basal ganglia |
| Caudate Cp L | 496 | -8 | 12 | 16 | 6.35 | Basal ganglia |
| Caudate Cp L | 248 | -12 | 18 | 14 | 4.17 | Basal ganglia |
| Bstem Med R | 320 | 2 | -38 | -46 | 3.62 | Brainstem |
| Bstem Ponscv R | 464 | 2 | -24 | -46 | 4.06 | Brainstem |
| Multiple regions | 11488 | -14 | -30 | -6 | 8.04 | Brainstem |
| Cblm CrusII R | 864 | 14 | -84 | -40 | 4.26 | Cerebellum |
| Cblm CrusI R | 264 | 44 | -50 | -36 | 3.75 | Cerebellum |
| Cblm CrusII R | 744 | 10 | -88 | -32 | 5.15 | Cerebellum |
| Cblm CrusI R | 408 | 44 | -78 | -30 | 3.97 | Cerebellum |
| Cblm VI R | 1808 | 34 | -46 | -24 | 5.04 | Cerebellum |
| Cblm CrusI L | 736 | -44 | -40 | -28 | 4.03 | Cerebellum |
| Cblm CrusII R | 568 | 16 | -90 | -26 | 4.06 | Cerebellum |
| Cblm I IV R | 208 | 10 | -34 | -14 | 3.59 | Cerebellum |
| Ctx TE1a R | 2008 | 60 | -14 | -16 | 5.1 | Cortex Default Mode A |
| Ctx 10d L | 192 | -20 | 66 | 14 | 3.04 | Cortex Default Mode A |
| Ctx 31pd R | 200 | 10 | -50 | 26 | 3.24 | Cortex Default Mode A |
| Ctx RSC R | 672 | 6 | -32 | 26 | 4.22 | Cortex Default Mode A |
| Ctx 7m L | 1464 | -8 | -56 | 30 | 3.56 | Cortex Default Mode A |
| Ctx PGs R | 2200 | 40 | -70 | 38 | 4.61 | Cortex Default Mode A |
| Ctx 8Ad L | 248 | -20 | 32 | 36 | 3.82 | Cortex Default Mode A |
| Ctx 8Ad R | 392 | 26 | 22 | 40 | 4.63 | Cortex Default Mode A |
| Ctx PGs L | 1064 | -44 | -68 | 42 | 4.15 | Cortex Default Mode A |
| Ctx 8Ad R | 272 | 14 | 34 | 42 | 3.96 | Cortex Default Mode A |
| Ctx TGd L | 6384 | -40 | 8 | -28 | 7.92 | Cortex Default Mode B |
| Ctx TE1a_L | 800 | -64 | -16 | -28 | 5.29 | Cortex Default Mode B |
| Ctx 47s L | 128 | -30 | 18 | -26 | 3.33 | Cortex Default Mode B |
| Ctx 45 L | 568 | -58 | 30 | 16 | 3.9 | Cortex Default Mode B |
| Ctx POS1 R | 192 | 8 | -52 | 14 | 3.06 | Cortex Default Mode C |
| Ctx PGi R | 160 | 40 | -58 | 28 | 3.36 | Cortex Default Mode C |
| Ctx RSC L | 824 | -6 | -16 | 30 | 5.62 | Cortex Default Mode C |
| Ctx TE2p L | 344 | -48 | -24 | -26 | 3.87 | Cortex Dorsal Attention A |
| Ctx FFC L | 400 | -50 | -60 | -16 | 3.73 | Cortex Dorsal Attention A |
| Ctx TE1m L | 176 | -54 | -28 | -18 | 3.9 | Cortex Fronto Parietal B |
| Ctx a10p R | 328 | 22 | 62 | -10 | 3.65 | Cortex Fronto Parietal B |
| Ctx a47r R | 576 | 30 | 62 | 2 | 3.96 | Cortex Fronto Parietal B |
| Ctx POS2 R | 440 | 0 | -70 | 26 | 3.2 | Cortex Fronto Parietal C |
| Ctx TGv R | 1016 | 42 | -6 | -42 | 4.01 | Cortex Limbic |
| Ctx TGd R | 472 | 24 | 16 | -40 | 4.88 | Cortex Limbic |
| Ctx TE2a L | 184 | -52 | -26 | -28 | 3.42 | Cortex Limbic |
| Ctx TGd R | 528 | 42 | 6 | -22 | 5.68 | Cortex Limbic |
| Ctx 13l L | 288 | -32 | 32 | -20 | 3.88 | Cortex Limbic |
| Ctx TF R | 192 | 38 | -26 | -14 | 3.37 | Cortex Limbic |
| Ctx 10pp R | 640 | 18 | 56 | -10 | 4.17 | Cortex Limbic |
| Ctx 3a L | 960 | -50 | -14 | 24 | 4.56 | Cortex Somatomotor A |
| Ctx 4 L | 152 | -50 | -6 | 24 | 3.33 | Cortex Somatomotor A |
| Ctx 1 L | 992 | -58 | -24 | 50 | 4.14 | Cortex Somatomotor A |
| Ctx 3b R | 152 | 46 | -14 | 42 | 3.48 | Cortex Somatomotor A |
| Ctx 6mp L | 536 | -16 | -22 | 64 | 3.83 | Cortex Somatomotor A |
| Ctx 6v L | 280 | -64 | 6 | 28 | 5.18 | Cortex Somatomotor B |
| Ctx STGa R | 432 | 52 | 8 | -22 | 3.32 | Cortex Temporal Parietal |
| Ctx A5 L | 2480 | -64 | -16 | -8 | 6.98 | Cortex Temporal Parietal |
| Ctx A5 L | 200 | -62 | 0 | -10 | 4.98 | Cortex Temporal Parietal |
| Ctx STSdp R | 1144 | 44 | -28 | 4 | 4.08 | Cortex Temporal Parietal |
| Ctx AAIC R | 1992 | 26 | 14 | -18 | 3.8 | Cortex Ventral Attention A |
| Ctx 6r L | 320 | -62 | 12 | 20 | 4.59 | Cortex Ventral Attention A |
| Ctx SCEF L | 472 | -4 | -2 | 50 | 3.63 | Cortex Ventral Attention A |
| Ctx SCEF R | 392 | 4 | 8 | 52 | 3.8 | Cortex Ventral Attention A |
| Ctx AVI L | 320 | -30 | 18 | -8 | 3.51 | Cortex Ventral Attention B |
| Ctx AVI L | 864 | -30 | 26 | -8 | 3.93 | Cortex Ventral Attention B |
| Ctx FOP5 R | 384 | 38 | 30 | -4 | 3.71 | Cortex Ventral Attention B |
| Ctx 46 L | 568 | -50 | 40 | 16 | 3.55 | Cortex Ventral Attention B |
| Ctx 9 46d L | 488 | -28 | 38 | 22 | 3.95 | Cortex Ventral Attention B |
| Ctx 46 R | 280 | 42 | 50 | 26 | 3.65 | Cortex Ventral Attention B |
| Ctx PIT L | 560 | -52 | -72 | -10 | 3.71 | Cortex Visual Central |
| Ctx V1 L | 1048 | -4 | -58 | 4 | 3.75 | Cortex Visual Peripheral |
| Ctx ProS L | 552 | -28 | -54 | 4 | 3.47 | Cortex Visual Peripheral |
| Ctx V6A R | 320 | 20 | -86 | 42 | 3.26 | Cortex Visual Peripheral |
| Thal Pulv | 5080 | 10 | -30 | 2 | 6.78 | Diencephalon |
| No label | 264 | 34 | -46 | 0 | 5.4 | No description |
| No label | 352 | 32 | -50 | 6 | 4.07 | No description |
| No label | 624 | 26 | -46 | 20 | 4.54 | No description |
| No label | 224 | 4 | 8 | 26 | 4.33 | No description |

**Supplementary Table 4.** Moderation of *Path ab* effects by individual differences in explicit learning (FDR corrected *q* < 0.05).

| Region | Volume<br>(mm3) | X | Y | Z | maxZ | Large-scale region or network |
| --- | --- | --- | --- | --- | --- | --- |
| **Moderation of *path ab* effects by individual differences in explicit learning** |  |  |  |  |  |  |
| CA3 Hippocampus | 304 | -32 | -22 | -12 | 3.62 | Hippocampus |

## Notes

### Competing Interest Statement

The authors have declared no competing interest.

